# Identifying Severe Stroke Patients Likely to Benefit From Thrombectomy Despite Delays of up to a Day

**DOI:** 10.1101/526947

**Authors:** R. Gilberto González, Gisele Sampaio Silva, Julian He, Saloomeh Sadaghiani, Ona Wu, Aneesh B. Singhal

## Abstract

Selected patients with large vessel occlusions (LVO) can benefit from thrombectomy up to 24 hours after onset. Identifying patients who might benefit from late intervention after transfer from community hospitals to thrombectomy-capable centers would be valuable. We searched for presentation biomarkers to identify such patients. Frequent MR imaging over 2 days of 38 untreated LVO patients revealed logarithmic growth of the ischemic infarct core. In 24 patients with terminal internal carotid artery or the proximal middle cerebral artery occlusions we found that an infarct core growth rate (IGR) <4.1 ml/hr and initial infarct core volumes (ICV) <19.9 ml had accuracies >89% for identifying patients who would still have a core of <50ml 24 hours after stroke onset, a core size that should predict favorable outcomes with thrombectomy. Published reports indicate that up to half of all LVO stroke patients have an IGR<4.1 ml/hr. Other potentially useful biomarkers include the NIHSS and the perfusion measurements MTT and Tmax. We conclude that many LVO patients have a stroke physiology that is favorable for late intervention, and that there are biomarkers that can accurately identify them at early time points as suitable for transfer for intervention.

## Introduction

Patients who suffer strokes caused by large vessel occlusions of the anterior circulation (LVOs, viz., terminal internal cerebral artery or middle cerebral artery M1/M2 occlusions) typically will have severe neurological deficits and poor outcomes if not treated. ^1^ While they comprise approximately a third of all ischemic strokes, they cause most deaths and poor outcomes. Clinical trials have shown that LVO-stroke patients may be effectively treated with thrombectomy within 6-8 hours.^2-8^ More recently, the DAWN^9^and DEFUSE-3^10^ trials have proven that thrombectomy may be successful up to 24 hours *post ictus* in carefully selected LVO patients. The widening of the thrombectomy time window presents new opportunities to treat LVO stroke patients; the challenge in community hospitals is to identify patients who will benefit from thrombectomy despite late arrivals or delays due to transfer to intervention-capable centers. We analyzed our cohort of LVO patients who underwent frequent MR imaging over 2 days to identify biomarkers on the admission scan that accurately identify patients likely to have favorable outcomes with thrombectomy despite sustaining significant delays.

Our aim was to identify patient biomarkers measured at first hospital arrival that would predict that the infarct core at 24 hours after stroke onset would be no larger than 50ml. The time and core size targets are conservative. Most patients in countries in North America, Europe and Japan are well within a 24-hour drive from a thrombectomy-capable center. The ischemic core target volume of 50ml was based on the data from the prospective thrombectomy clinical trials that used core volumes of less 50ml ^6,9^ as inclusion criteria, as well as the observation from the HERMES collaboration^11^ that revealed that patients with final infarcts of <50ml after thrombectomy had an ∼80% probability of living independently at 3 months even when they presented with severe neurological deficits.

## Methods

### Patient selection and evaluation

This study was compliant with the Health Insurance Portability and Accountability Act (HIPAA) and was approved by Partners Human Research Committee / Massachusetts General Hospital institutional review board (IRB). The data are from the Normobaric Oxygen Therapy (NBO) in Acute Ischemic Stroke Trial, in which all methods were carried out in accordance with relevant guidelines and regulations and that informed consent was obtained from all participants or, if participants are under 18, from a parent and/or legal guardian. (see http://clinicaltrials.gov/show/NCT00414726 for the complete trial inclusion and exclusion criteria). We included patients who met the following criteria: 1) MR imaging including a DWI scan showing acute ischemic injury; 2) CT angiography (CTA) or MR angiography (MRA) of the head demonstrating a proximal anterior circulation artery occlusion (i.e. terminal ICA and/or proximal MCA (M1 and/or M2 origin); 3) three or more MRIs performed within ∼2 days of stroke onset. Of the 60 subjects who underwent serial MRI in the clinical trial, 38 met inclusion criteria for this analysis. For patients whose stroke onset was not witnessed, onset time was estimated as midway between last seen well and first seen with stroke symptoms. ^21^ National Institutes of Health Stroke Scale (NIHSS) scores were recorded at all time points, and the modified Rankin Scale (mRS) was recorded 3 months after admission, by investigators blinded to MRI lesions and treatment assignment. A favorable outcome was defined as living independently (mRS score of 2 or less) at 3 months.

### Imaging Techniques

MRI scans were obtained at admission and repeated after ∼12 hours, ∼1 day and ∼2 days after onset using a clinical 1.5-T (General Electric, Waukesha, Wisconsin) MRI system. Sagittal T1, axial DWI, Fluid-attenuated inversion recovery (FLAIR) T2, and gradient-echo sequences were performed at each time point. In addition, head MRA was obtained with the first three MRI scans.

DWI were acquired using the following median values: field of view of 220 mm, 25 slices, thickness of 5 mm, gap of 1 mm, TR of 5 seconds, TE of 85.3 ms, acquisition matrix 128×128, and with b=0 s/mm2 and b=1000 s/mm2 in at least 6 diffusion-gradient directions. Isotropic DWI and apparent diffusion coefficient (ADC) maps were calculated using techniques previously described.^22^ FLAIR imaging were performed with a fast-spin echo sequence, with the following median values: 10s TR, 145ms TE, 2200ms TI, 256×192 matrix, 220mm FOV, 25 5mm slices with 1mm gap. Gradient-echo T2* imaging median values included TR/TE of 817/25ms, 20° flip angle, 256×192 matrix, 220 mm FOV, 25 5mm thick slices and 1 mm gap. Perfusion MRI data were acquired using the following median values: field of view of 220 mm, 15 slices, thickness of 5 mm, gap of 1 mm, TR of 1.5 seconds, TE of 40 ms, flip angle 60 degrees, acquisition matrix 128×128. 80 data points were acquired using gradient-echo echo planar imaging readout. Mean-transit time (MTT) and Tmax (time at which the tissue response function reached maximum value) perfusion maps were calculated using automated oscillation index regularized deconvolution. ^23^

The 3D time-of-flight MRA consisted of a 7cm-thick slab positioned over the circle of Willis. The median imaging parameters were TR/TE=36/6.8 ms, 25° flip-angle, 180 mm FOV, 320×192 matrix, 101 axial images, 1.4-mm-thick with 0.7-mm overlap. The MRA source images were post-processed into maximum intensity projection images.

CT angiography was performed using multi-detector scanners (GE Medical Systems, Milwaukee, WI) from the vertex to the aortic arch following injection of 65–140 ml of a nonionic contrast agent (Isovue; Bracco Diagnostics, Princeton, NJ) at a rate of 3 to 4 ml/s. The median parameters were 1.25-mm slice thickness, 220 mm reconstruction diameter, 120 kV, and 657 mA.

### Image Analysis

DWI abnormalities were outlined visually using both the DWI and ADC maps from the same time-point. Tmax lesions were outlined visually with knowledge of DWI and ADC, while MTT abnormalities were outlined visually with knowledge of DWI, ADC and Tmax maps. Analysts blinded to treatment assignment, time-point and clinical information performed all outlines. All outlines were performed using semi-automated, open-source software (Display, Montreal Neurologic Institute, Montreal). Lesion growth was calculated as the Infarct core volume divided by the time in hours from last seen well or from the prior measurement.

### Statistical Analysis

All values are reported as percentage, mean (standard error), or median. Mann-Whitney, Kruskal-Wallis, ANOVA, or Friedman Test, as appropriate, was used to assess changes in lesion volumes across the 4 time points. Correlation analyses were performed between DWI volumes and time as well as between DWI volumes and natural logarithm of time. Simple and multivariable regression analyses were used to assess the relationship of clinical and imaging variables measured at the time of admission with functional outcomes at 3 months. P-values of less than 0.05 were considered statistically significant. Receiver operator characteristic (ROC) analyses were performed to evaluate the sensitivity and specificity of presenting variables. Analyses described were performed using MedCalc version 14.8.1 and SPSS statistical software (release 20.0 for Windows; SPSS, Chicago, IL). ROC analyses were performed using MedCalc.

## Results

MR imaging was performed up to 3 times in the first day and once more at 2 days *post ictus* in 38 patients with acute anterior circulation LVO ischemic stroke. No patient received thrombolytic therapy or thrombectomy; 18 were treated with normobaric oxygen and 20 with medical air for 8 hours as part of a randomized clinical trial. Demographic and clinical information is displayed in Table 1. Initial MRI scans (MRI-1) were obtained at a mean time after stroke onset of 5.0 hours (range, 1.6-9.5 hours). Subsequent scans (MRI-2, MRI-3 and MRI-4) were acquired at a mean of 11.7 hours (range, 7-15 hours), 1 day (mean 25.8 hours; range, 22-33 hours), and 2 days (mean 49.7 hours; range, 43-57 hours) following stroke onset.

**Table 1.** LVO (N=38) Baseline Variables and Stroke Etiology.

|  |  |  |  |
| --- | --- | --- | --- |
| Age (yr) | 72 | Site of arterial occlusion |  |
| Sex |  | ICA | 8 |
| Female | 16 | M1 | 17 |
| Male | 22 | M2 | 13 |
| NIHSS | 14 | none of the above | 0 |
| Glucose (mg/dl) | 137 | 24-hour reperfusion |  |
| Treatment type |  | complete | 3 |
| NBO | 18 | partial | 14 |
| Medical air | 20 | none | 19 |
| Causative Classification System |  | unavailable | 2 |
| Large artery atherosclerosis | 5 | 24-hour recanalization |  |
| Cardio-aortic embolism | 23 | complete | 5 |
| Small artery occlusion | 0 | partial | 8 |
| Other uncommon causes | 5 | none | 22 |
| Undetermined causes | 5 | unavailable | 3 |

LVO: large vessel occlusion. NBO: Normobaric Oxygen Therapy; NIHSS: National Institutes of Health Stroke Scale; ICA, internal carotid artery; M1 and M2, segments of the middle cerebral artery

To illustrate the primary imaging data that was used for this study, Figure 1 shows selected diffusion and perfusion images from a patient with an occlusion of the M1 segment of the left middle cerebral artery. The patient had an initial core volume (ICV) of <50ml, as revealed by diffusion weighted imaging (DWI), and a much larger volume of abnormal cerebral perfusion. Notably, despite persistence of arterial occlusion and the large perfusion defect, the ischemic core grew slowly over 48 hours and remained <50ml.

**Figure 1.**
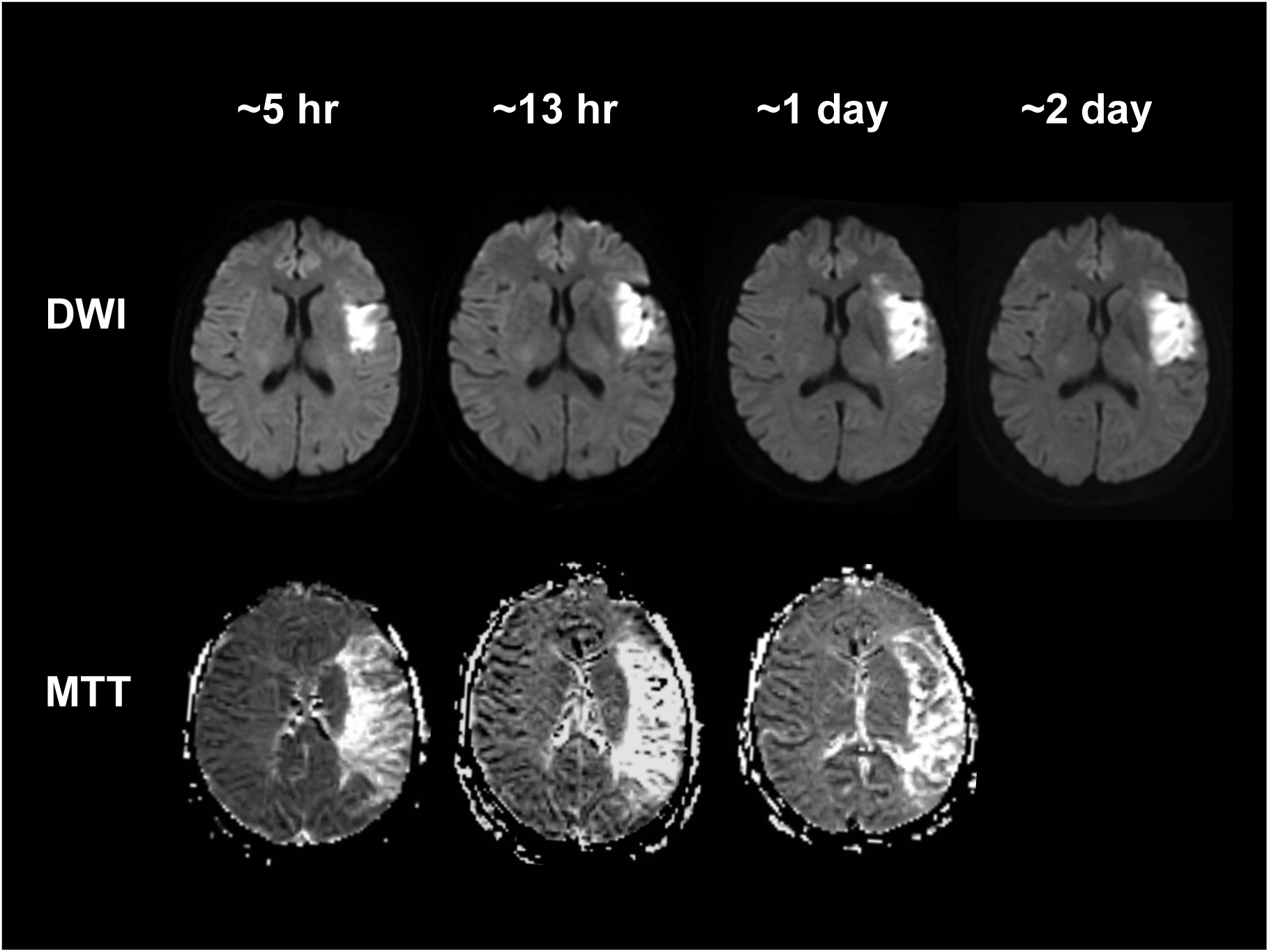
Diffusion and perfusion mean transit time images of patient with a left middle cerebral artery mainstem occlusion that persisted at 24 hours. Diffusion and perfusion MRI studies were performed at ∼5 hours, ∼ 13 hours, ∼1 day and ∼2 days from last seen well.

### Ischemic core volume growth rates in patients with ICA, M1 & M2 occlusions

Table 2 displays the average ischemic lesion growth in LVO patients grouped by initial ICV. The highest average growth occurred during the initial period between stroke onset and the time of the first imaging. In all groups, the second period (∼5-12 hours) the average growth of the core was less than 10% of the initial period. The lesion growth during the other periods was also much lower than the initial period for all groups. Infarct core growth is nonlinear: linear regression resulted in a poor correlation coefficient between lesion volumes and time (R^2^<0.6); conversely, correlation coefficients were high (R^2^≥0.96) between lesion volumes and the natural logarithm of time, suggesting logarithmic growth of the ischemic core over 2 days.

**Table 2.** Diffusion Lesion Growth Rates.

| Group | MRI-1<br>lesion<br>volume<br>ml(SE) | Symptom<br>Onset to MRI-1<br>growth rate<br>ml/hr(SE) | MRI-1 to<br>MRI-2<br>growth rate<br>ml/hr(SE) | MRI-2 to<br>MRI-3<br>growth rate<br>ml/hr(SE) | MRI-3 to<br>MRI-4<br>growth rate<br>ml/hr(SE) | P<br>value* |
| --- | --- | --- | --- | --- | --- | --- |
| *LVO patients grouped by admission infarct core volume* |  |  |  |  |  |  |
| <50 mL<br>(n=23) | 20 (3.5) | 4.0 (0.8) | 0.4 (0.3) | 0.6 (0.2) | 0.4 (0.1) | <0.01 |
| 50-100 mL<br>(n=6) | 66 (1.7) | 12.5 (1.4) | 1.2 (1.3) | 3.3 (1.5) | 1 (0.3) | 0.08 |
| >100 mL<br>(n=9) | 139(14.5) | 29.5 (1.9) | 1.8 (1.6) | 4.3 (1.1) | 2 (0.8) | <0.01 |
| *P value\** | <0.01 | <0.01 | 0.74 | <0.01 | <0.01 |  |
| *LVO patients with persistent occlusions at 1 day grouped by admission infarct core volume* |  |  |  |  |  |  |
| <50mL<br>(n=14) | 15 (3.5) | 3.4 (1.1) | 0.1 (0.2) | 0.8 (0.3) | 0.5 (0.1) | <0.01 |
| 50-100 mL<br>(n=3) | 66 (2.4) | 13.3 (2.7) | 2.4 (1.5) | 5.3 (2.6) | 1.4 (0.3) | 0.24 |
| >100 mL<br>(n=5) | 138 (8.7) | 28.1 (2.6) | 2.6 (2.8) | 5.5 (1.9) | 2.8 (1.2) | 0.01 |
| *P value\** | <0.01 | <0.01 | 0.38 | <0.01 | <0.01 |  |

\*Mann-Whitney, Kuskal-Wallis, ANOVA, or Friedman Test, as appropriate.
Abbreviations: NBO, normobaric oxygen therapy; ICA, internal carotid artery; MCA, middle cerebral artery; DWI, diffusion lesion volume. MRI-1 performed at mean 5.0 hours (range, 1.6-9.5 hours) after stroke onset; MRI-2, 11.7 hours (range, 7-15 hours); MRI-3, 25.8 hours (range, 22-33 hours); and MRI-4, 49.7 hours (range, 43-57 hours) after stroke onset.

### Temporal Ischemic Core Volume Changes in Patients with ICA and M1 Occlusions

Figure 2 displays the changes in ischemic core volumes, grouped by baseline infarct core volumes in the 7 patients with distal ICA and 17 patients with M1 occlusions. Patients with small initial infarct cores (<50ml) at baseline had slower core growth compared to patients with large baseline cores (>100ml).

**Figure 2.**
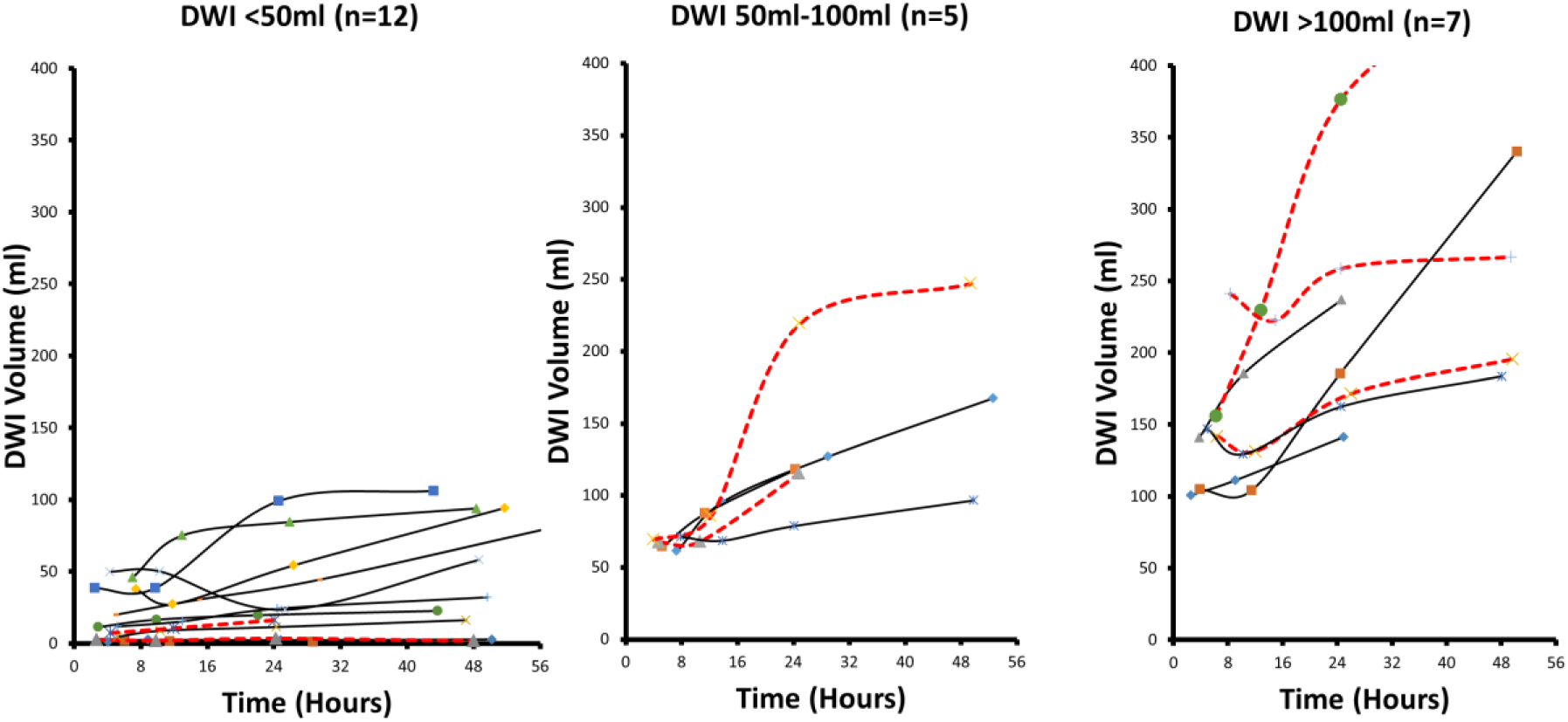
Ischemic core growth in 24 patients with M1 (solid line) and ICA (dashed lines) occlusions. Left. 12 patients with baseline infarct core volume less than 50 ml. Middle. 5 patients with an ICV of between 50 and 100 ml. Right. 7 patients with an ICV greater than 100 ml.

### Biomarkers of Slow Progressors Likely to Benefit from Delayed Thrombectomy

Our goal was to identify the biomarkers that identify patients that have such slow growth of infarct cores that they may be amenable to successful thrombectomy up to 1 day after onset. Other researchers, most notably the Pittsburgh group ^12^, have emphasized the importance of slow progressors and their management. Here we adopt a practical definition of slow progressors as patients with an infarct core of appropriate size to be amenable to successful thrombectomy up to 1 day after onset. We used a target ischemic core size of <50ml at the time of the 1-day scan to identify these patients. This conservative core target of 50ml was based on the data from the prospective clinical trials that used core volumes of less than 50ml ^6,9^ as inclusion criteria, as well as the observation from the HERMES collaboration^11^ that patients with final infarcts of <50ml after thrombectomy had an ∼80% probability of living independently at 3 months irrespective of initial symptom severity.

Receiver operator characteristic (ROC) analyses were performed to find the best biomarkers of treatable slow progressors in the 24 patients with ICA and M1 occlusions. We compared key characteristics of our patient group with the *control* cohorts from the DEFUSE3 and DAWN thrombectomy clinical trials and is shown in Table 3.

**Table 3.** Comparison of patient characteristics of ICA/M1 occlusion patients with DEFUSE-3^10^ and DAWN ^9^control groups

|  | Our cohort<br>(N=24) | DEFUSE-3 Controls<br>(N=90) | DAWN Controls<br>(N=99) |
| --- | --- | --- | --- |
| Age | 70 (median) | 71 (median) | 71 (mean) |
| Females | 33% | 51% | 43% |
| Median NIHSS | 17 | 16 | 17 |
| M1 occlusions | 68% | 60% | 78% |
| Favorable outcomes | 13% | 17% | 13% |

All of the variables are similar to trial control groups with the exception of a lower proportion of female patients in our cohort. Admission variables that were evaluated included NIHSS, admission infarct core volume (ICV), initial infarct core growth rate (IGR), mean transit time volume, Tmax volume, and the diffusion/perfusion mismatch. Figure 3 displays an ROC graph showing the sensitivity versus the false positive rate for predicting the <50ml target volume at 24 hours for 3 of these variables at the time of presentation.

**Figure 3.**
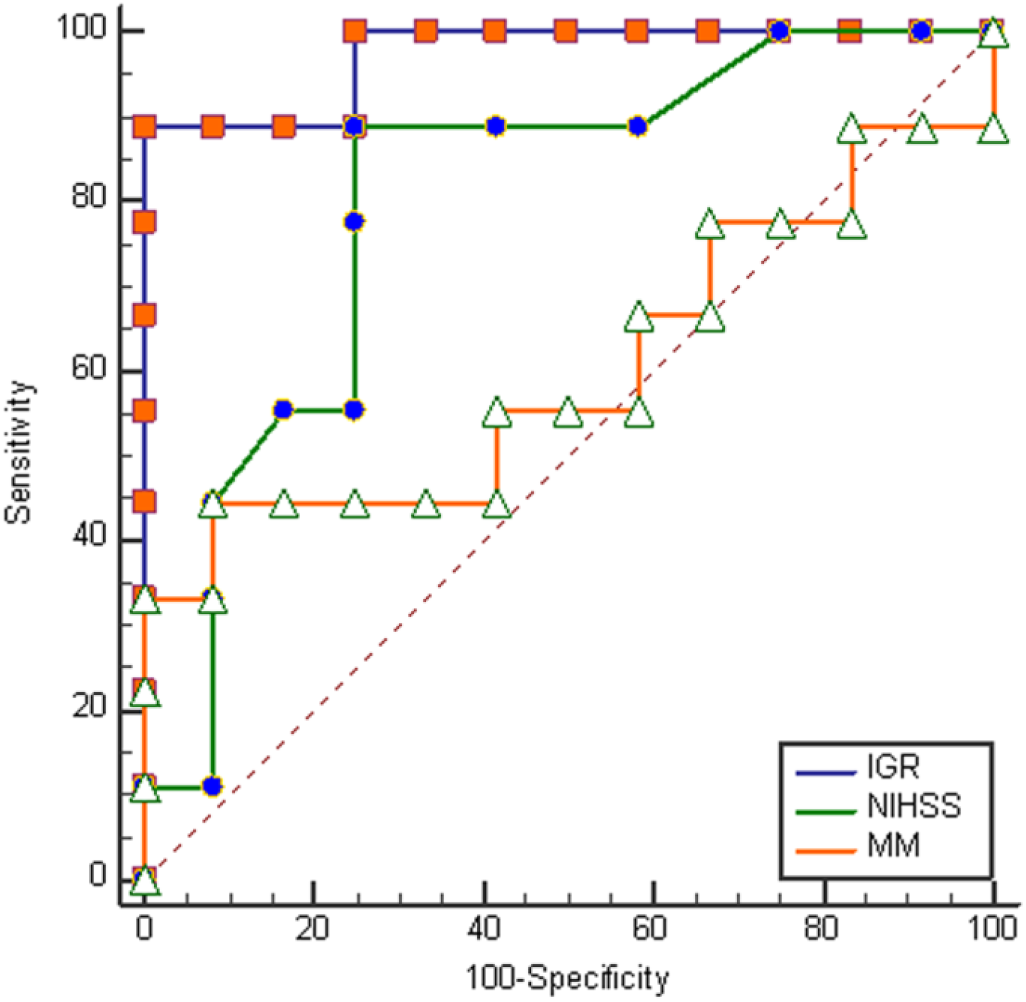
Receiver operator characteristic (ROC) curves of variables predicting ischemic core volumes <50 ml one day after stroke onset. Data is from the 24 patients with an ICA or M1 occlusion identified on CT or MR angiography. For clarity of presentation, the only variables included in this graph ischemic stroke initial growth rate (IGR), NIH Stroke Scale (NIHSS), and diffusion/perfusion mismatch (MM).

More details of the ROC analyses are shown in Table 4. IGR and ICV were equivalent as the best performing markers with the highest sensitivities and the lowest false positive rates. Both had areas under the curve (AUC) of 0.97 or better as well as 89% sensitivity and 100% specificity for the specific criteria of <4.1 ml/hr (IGR) and <19.9 ml (ICV).

**Table 4.** ROC Analyses of Baseline Variables To Predict Ischemic Core of < 50ml One Day *Post Ictus*

| Variable | Criteria | Sensitivity | Specificity | AUC (95% CI) |
| --- | --- | --- | --- | --- |
| Infarct Growth Rate | <4.1 ml/hr | 89% | 100% | 0.97 (0.81-1.0) |
| Initial Core Volume | <19.9 ml | 89% | 100% | 0.98 (0.82-1.0) |
| MTT | <150 ml | 89% | 83% | 0.85 (0.63-0.97) |
| NIHSS | ≤16 | 89% | 73% | 0.80 (0.59-0.94) |
| Tmax | <121 ml | 56% | 100% | 0.75 (0.52-0.91) |
| Diff/Perf Mismatch | <43 ml | 44% | 92% | 0.60 (0.37-0.81) |

## Discussion

Patients with potentially severe strokes caused by an occlusion of a major cerebral artery but with a favorable physiology have small infarct cores that grow slowly and are likely to be excellent candidates for thrombectomy despite significant time delays. This finding comes from a cohort of LVO patients who were part of a clinical trial that assessed the value of normobaric oxygen (NBO). While NBO was not shown to be effective ^13^, the study design included multiple MRI scans over a short time and resulted in the unique observations described here. Using a target ischemic core volume of 50 ml at 24 hours, we found that that the best biomarkers of such a favorable physiology are an infarct core growth rate of <4.1 ml/hr or an initial infarct core volume of <19.9 ml.

We chose 50 ml target volume after considering a report from the HERMES collaboration (that combined data from 7 thrombectomy trials) that indicated a 50 ml final infarct volume *after* thrombectomy results in a high probability of excellent long-term outcomes despite the severity of presenting symptoms. ^11^ Our targets are not meant to be prescriptive. Other target volumes may be better. For example, ROC analysis of a 70 ml target revealed 90%/100% sensitivity/specificity for an IGR <5.2 ml. Suitable volumes may be even larger if there is less eloquence of the tissue at risk, and investigations of larger target volumes in patients with lower NIHSS may also be very useful. The assessment at 24 hours is also somewhat arbitrary, and was only used because that is the outer time limit of the successful trials published to date. It is very likely that there will be patients that will benefit from treatment beyond 24 hours.

The reliable identification of LVO slow progressors could greatly expand the use of thrombectomy, and more widely apply the lessons from the recent DAWN ^9^ and DEFUSE-3^10^ thrombectomy trials that treated patients up to 24 hours after onset. The substantial number of slow progressors is largely explained by collateral physiology that typically produces logarithmic growth of the ischemic core as documented in this study. In animal stroke models, the ischemic core also grows in a logarithmic fashion. ^14^ Documentation of this type of growth in patients has lacked empirical validation. Wheeler et al. ^15^ suggested logarithmic infarct growth rates in LVO patients using only 2 imaging data points: baseline and 1-week core volumes. Because 4 imaging studies were performed over 2 days, we are able to validate logarithmic-type growth, and explain on our prior observation of apparent ischemic core stability in LVO patients.^16^ We found that most of the core growth occurred within the first 6 hours, with less than 10% further growth over remainder of the first day. Comparison of line fits of core volume with respect to time or the natural logarithm of time favored logarithmic growth.

Our findings beg the question of what proportion of all LVO stroke patients are slow progressors. A review of published studies of patients with LVO that included IGR measurements suggests that a large proportion have growth rates below 4.1 ml/hr and are slow progressors as we have defined them here (Table 5). Taken together, the published evidence from nearly 700 LVO patients suggest that up to half of LVO patients are slow progressors. If this is confirmed, the targeted thrombectomy of slow progressors has the potential to substantially reduce the deaths, severe disabilities and costs due to stroke.

**Table 5.** Studies of large vessel occlusion stroke patients with published initial infarct growth rates (IGR). ^19, 15, 24, 20^,^25^

| Median IGR (ml/hr) | Sample size | References |
| --- | --- | --- |
| 4.2 | 186 | Hakimelahi R et al. |
| 3.1 | 65 | Wheeler et al. |
| 6.1 | 166 | Olivot et al. |
| 5.2 | 94 | Vagal et al. |
| 2.6 | 185 | Desai, et al. |

The very high variability of lesion growth in LVO patients is best explained by differences in the collateral circulation. ^17-20^ Slow progressors must have robust collateral circulations that persist for long periods. An important consideration is whether slow progressor LVO patients will still need thrombectomy after a long delay or will spontaneous recanalization make the procedure unnecessary. We found that only 13% of slow progressors with ICA or M1 occlusions had favorable functional outcomes at 3 months, similar to the control cohorts in the DAWN and DEFUSE-3 thrombectomy clinical trials. Not undergoing thrombectomy for such occlusions is clearly hazardous, even in slow progressors, because the arterial occlusion may persist for a long period and the advantageous collateral physiology may diminish and ultimately collapse.

There may be more practical but still effective alternatives to MRI for the identification of LVO patients that are slow progressors. Our ROC analyses (Table 4) hint at several approaches, and there may be others. This is important because patients may present at centers without an available MRI. One alternative may be CT perfusion. We found that the MR perfusion measurement MTT performed well, and it is likely that a CT perfusion MTT measurement would be similarly effective. Additionally, while not tested in this study, perfusion CT estimates of the ischemic core may be effective. A marker that does not rely on imaging is the NIHSS score of <16 in patients with ICA/M1 occlusions: about 90% of slow progressors would be identified, with a false positive rate would be 3 out of 10. A different NIHSS target might be chosen for a higher specificity, but with a lower sensitivity. For example, an NIHSS ≤9 is 100% specific and 63% sensitive. There are likely other effective imaging biomarkers that may derived from CT perfusion, CT angiography, and possibly even noncontrast CT data.

An important strength of this study is that it documented the natural history of ischemic core evolution, without confounders such as thrombolytic use or thrombectomy, which has rarely been done previously.^20^ A possible limitation is the relatively small number of patients that were studied. However, the fact that each patient was scanned multiple times makes our findings robust. A cross sectional study designed to reveal the findings described here would likely require several hundred patients. There is the potential confound of NBO therapy on the NBO-treated arm; however, there was no significant difference between the NBO and room air treated groups on any imaging variable that was measured including outcomes. ^13^ In any case, prospective studies are needed to determine the value of imaging biomarkers in making triage decisions for transport of patients for thrombectomy.

In conclusion, ischemic core initial growth rates of <4.1 ml/hr and ischemic core volumes of <19.9 ml are excellent biomarkers for selecting LVO patients that are likely to benefit from transfer to thrombectomy-capable stroke centers despite a substantial delay. These biomarkers are effective because of the characteristic physiological logarithmic growth of the ischemic core. Published accounts suggest that up to half of all LVO stroke patients meet these criteria. Other biomarkers based on CT imaging or NIHSS may be more practical and more widely applicable, although they are somewhat less effective. These observations herald an opportunity to treat many more severe ischemic stroke patients with thrombectomy by transporting patients identified with suitable biomarkers to have a favorable physiology.

## Data Availability

The datasets generated during and/or analyzed during the current study are available from the corresponding author on reasonable request.

## Acknowledgements

The following founding sources are provided by authors, NIH-NINDS R01NS051412 (ABS, GSS, SS, OW), P50NS051343 (ABS, GSS, SS, OW), R21NS077442 (ABS), R21-NS085574 (ABS), U10 -NS086729 (ABS), R01NS059775 (OW), and R01NS063925 (OW). Current address for Dr. Gisele S Silva is Dept. of Neurology, Hospital Israelita Albert Einstein, São Paulo, Brazil.

## Author Contributions

ABS and RGG conceived and designed the study. RGG wrote the first draft. GSS, JH, SS, and OW did the analysis. All authors contributed to the interpretation of the results. ABS is integrity guarantor.

## References

1 Gonzalez, R. G. et al. Good outcome rate of 35% in IV-tPA-treated patients with computed tomography angiography confirmed severe anterior circulation occlusive stroke. Stroke; a journal of cerebral circulation 44, 3109–3113, doi: 10.1161/STROKEAHA.113.001938 (2013).

2 Berkhemer, O. A. et al. A randomized trial of intraarterial treatment for acute ischemic stroke. The New England journal of medicine 372, 11–20, doi: 10.1056/NEJMoa1411587 (2015).

3 Campbell, B. C. et al. Endovascular therapy for ischemic stroke with perfusion-imaging selection. The New England journal of medicine 372, 1009–1018, doi: 10.1056/NEJMoa1414792 (2015).

4 Goyal, M. et al. Randomized assessment of rapid endovascular treatment of ischemic stroke. The New England journal of medicine 372, 1019–1030, doi: 10.1056/NEJMoa1414905 (2015).

5 Jovin, T. G. et al. Thrombectomy within 8 hours after symptom onset in ischemic stroke. The New England journal of medicine 372, 2296–2306, doi: 10.1056/NEJMoa1503780 (2015).

6 Saver, J. L. et al. Stent-retriever thrombectomy after intravenous t-PA vs. t-PA alone in stroke. The New England journal of medicine 372, 2285–2295, doi: 10.1056/NEJMoa1415061 (2015).

7 Bracard, S. et al. Mechanical thrombectomy after intravenous alteplase versus alteplase alone after stroke (THRACE): a randomised controlled trial. Lancet Neurol 15, 1138–1147, doi: 10.1016/S1474-4422(16)30177-6 (2016).

8 Muir, K. W. et al. Endovascular therapy for acute ischaemic stroke: the Pragmatic Ischaemic Stroke Thrombectomy Evaluation (PISTE) randomised, controlled trial. J Neurol Neurosurg Psychiatry 88, 38–44, doi: 10.1136/jnnp-2016-314117 (2017).

9 Nogueira, R. G. et al. Thrombectomy 6 to 24 Hours after Stroke with a Mismatch between Deficit and Infarct. The New England journal of medicine 378, 11–21, doi: 10.1056/NEJMoa1706442 (2018).

10 Albers, G. W. et al. Thrombectomy for Stroke at 6 to 16 Hours with Selection by Perfusion Imaging. The New England journal of medicine 378, 708–718, doi: 10.1056/NEJMoa1713973 (2018).

11 Boers, A. M. M. et al. Association of follow-up infarct volume with functional outcome in acute ischemic stroke: a pooled analysis of seven randomized trials. J Neurointerv Surg 10, 1137–1142, doi: 10.1136/neurintsurg-2017-013724 (2018).

12 Rocha, M. & Jovin, T. G. Fast Versus Slow Progressors of Infarct Growth in Large Vessel Occlusion Stroke: Clinical and Research Implications. Stroke; a journal of cerebral circulation 48, 2621–2627, doi: 10.1161/STROKEAHA.117.017673 (2017).

13 Singhal, A. & Investigators, P. S. Serial MRI Findings in a Phase 2B Clinical Trial of Normobaric Oxygen Therapy (NBO) in Acute Ischemic Stroke (AIS). Neurology 80(suppl), S42.001 (2013).

14 Bardutzky, J. et al. Perfusion and diffusion imaging in acute focal cerebral ischemia: temporal vs. spatial resolution. Brain research 1043, 155–162, doi: 10.1016/j.brainres.2005.02.073 (2005).

15 Wheeler, H. M. et al. The growth rate of early DWI lesions is highly variable and associated with penumbral salvage and clinical outcomes following endovascular reperfusion. International journal of stroke: official journal of the International Stroke Society 10, 723–729, doi: 10.1111/ijs.12436 (2015).

16 Gonzalez, R. G. et al. Stability of large diffusion/perfusion mismatch in anterior circulation strokes for 4 or more hours. BMC neurology 10, 13, doi: 10.1186/1471-2377-10-13 (2010).

17 Christoforidis, G. A., Mohammad, Y., Kehagias, D., Avutu, B. & Slivka, A. P. Angiographic assessment of pial collaterals as a prognostic indicator following intra-arterial thrombolysis for acute ischemic stroke. AJNR Am J Neuroradiol 26, 1789–1797, doi: http://www.ajnr.org/content/26/7/1789.long (2005).

18 Liebeskind, D. S. Collaterals in acute stroke: beyond the clot. Neuroimaging Clin N Am 15, 553-573, x, doi: https://doi.org/10.1016/j.nic.2005.08.012 (2005).

19 Hakimelahi, R. et al. Time and diffusion lesion size in major anterior circulation ischemic strokes. Stroke; a journal of cerebral circulation 45, 2936–2941, doi: 10.1161/STROKEAHA.114.005644 (2014).

20 Vagal, A. et al. Collateral Clock Is More Important Than Time Clock for Tissue Fate. Stroke; a journal of cerebral circulation 49, 2102–2107, doi: 10.1161/STROKEAHA.118.021484 (2018).

21 Maas, M. B. & Singhal, A. B. Unwitnessed stroke: impact of different onset times on eligibility into stroke trials. Journal of stroke and cerebrovascular diseases: the official journal of National Stroke Association 22, 241–243, doi: 10.1016/j.jstrokecerebrovasdis.2011.08.004 (2013).

22 Sorensen, A. G. et al. Human acute cerebral ischemia: detection of changes in water diffusion anisotropy by using MR imaging. Radiology 212, 785–792, doi: 10.1148/radiology.212.3.r99se24785 (1999).

23 Copen, W. A. et al. Exposing hidden truncation-related errors in acute stroke perfusion imaging. AJNR Am J Neuroradiol 36, 638–645, doi: 10.3174/ajnr.A4186 (2015).

24 Olivot, J. M. et al. Impact of Initial Diffusion-Weighted Imaging Lesion Growth Rate on the Success of Endovascular Reperfusion Therapy. Stroke; a journal of cerebral circulation 47, 2305–2310, doi: 10.1161/STROKEAHA.116.013916 (2016).

25 Desai, S. M., Rocha, M., Jovin, T. G. & Jadhav, A. P. High Variability in Neuronal Loss Time Is Brain, Requantified. Stroke; a journal of cerebral circulation 50, 34–37 (2019).

